# The same ultra-rapid parallel brain dynamics underpin the production and perception of speech

**DOI:** 10.1101/2021.02.11.430723

**Authors:** Amie Fairs, Amandine Michelas, Sophie Dufour, Kristof Strijkers

## Abstract

The temporal dynamics by which linguistic information becomes available is one of the key properties to understand how language is organised in the brain. An unresolved debate between different brain language models is whether words, the building blocks of language, are activated in a sequential or parallel manner. In this study we approached this issue from a novel perspective by directly comparing the time course of word component activation in speech production versus perception. In an overt object naming task and a passive listening task we analysed with mixed linear models at the single-trial level the event-related brain potentials elicited by the same lexico-semantic and phonological word knowledge in the two language modalities. Results revealed that both word components manifested simultaneously as early as 75 ms after stimulus onset in production and perception; differences between the language modalities only became apparent after 300 ms of processing. The data provide evidence for ultra-rapid parallel dynamics of language processing and are interpreted within a neural assembly framework where words recruit the same integrated cell assemblies across production and perception. These word assemblies ignite early on in parallel and only later on reverberate in a behaviour-specific manner.

---

Language behaviour concerns a complex, multi-component process where different linguistic representations (semantic, lexical, syntactic and phonological knowledge) need to be retrieved and merged together in order to produce and perceive communicative signals. Insights on ‘when’ and ‘how’ different linguistic knowledge becomes activated is therefore key to understand the cortical mechanics that can sustain this uniquely human behaviour (e.g., Friederici, 2002; 2011; Indefrey & Levelt, 2004; Pulvermuller, 1999; 2018; Hauk, 2016). Despite the time-course of language processing being a longstanding question in the field, debate remains between proponents of more sequential dynamics versus those arguing for parallel retrieval of linguistic knowledge. At least two main reasons exist for this debate: first, in terms of data, empirical evidence has been found for both sequential and parallel brain dynamics; second, it is often difficult to disentangle whether neurophysiological responses to linguistic knowledge support a sequential or parallel time-course. This is in part due to the fact that language processing recruits a vast number of regions in the brain, making it hard to know whether temporal differences reflect functionally distinct mental operations or merely physical distance in the brain (and different axonal conductance velocities) but functionally parallel activations. Interestingly though, the debate concerning the temporal dynamics in brain language models exists both for speech production and speech perception, even though traditionally these modalities have been studied separately (e.g., Price, 2012). This shared debate (but separated approach) is interesting because directly contrasting the time-course of language processing between the production and perception of words could circumvent the above-mentioned conceptual problems and provide novel and more explicit insights on how our brain computes language in time. This is because a sequential model predicts the reverse temporal dynamics between the production and perception of words, while a parallel model predicts the same, simultaneous onset of word components across the language modalities. Therefore, in the current study, we systematically contrasted the event-related brain potentials (ERPs) elicited at the single-trial level by lexico-semantic and phonological word knowledge in both production and perception.

In sequential models of word production (e.g., Levelt et al., 1999; Indefrey & Levelt, 2004; Indefrey, 2011; Hickok, 2012) and perception (e.g., Friederici, 2002; 2011; Scott & Johnsrude, 2003; Hickok & Poeppel, 2007) the idea is that different word components are functionally separated in time (and space): In production first the semantic and lexical information of to-be-uttered words are retrieved, followed by the phonological code, which is then translated into a motor program to trigger the vocal folds for articulation. In perception, incoming acoustic speech signals are phonologically encoded and subsequently activate the lexical and semantic knowledge associated with those phonological forms. Though sequential models differ in how many processing stages are necessary to go from concept to speech or speech to concept, and can incorporate cascading and interactivity between processing layers (e.g., Dell, 1986; Caramazza, 1997; Friederici, 2011) rather than being strictly serial (e.g., Levelt et al., 1999; Friederici, 2002; Indefrey & Levelt, 2004), they do share the common feature that upstream processes should initiate prior to downstream processes within the hierarchical architecture (e.g., Dell & O’Sheaghda, 1992; Indefrey & Levelt, 2004; Friederici, 2011). Neurophysiological evidence supporting such sequential dynamics show that in speech production lexico-semantic effects emerge earlier in time (roughly around 200-250ms) than phonological and phonetic effects (roughly around 300-400ms) during speech production tasks (e.g., Salmelin et al., 1994; Levelt et al., 1998; Van Turennout et al., 1998; Vihla et al., 2006; Laganaro et al., 2009; Sahin et al., 2009; Schuhmann et al., 2012; Valente et al., 2014; Dubarry et al., 2017). Similar sequential-like neurophysiological responses have been found in speech perception (but in the reverse direction), with phonetic and phonological effects emerging roughly within the first 200 ms of processing and lexico-semantic effects manifesting roughly around 300-400 ms after spoken word onset (e.g., Holcomb & Neville, 1990;; Friederici et al.,1993; Van Petten et al., 1999; Hagoort & Brown, 2000; Halgren et al., 2002; O’Rourke & Holcomb, 2002; Dufour et al., 2013; Winsler et al., 2018).

The above temporal evidence paints a mirror image between language behaviours where cortical activation of different word components seems to progress in functional delays of some 100 ms and with reversed order between speech production and perception. However, this temporal segregation between the retrieval of meaning and sounds for words has received criticisms both in language perception (e.g., Pulvermuller et al., 2009) and production (e.g., Strijkers & Costa, 2011), and inconsistent evidence has been presented in both domains of language. In language perception, electrophysiological and neuromagnetic data have demonstrated that semantic, lexical and phonological word properties can emerge very rapidly (within 200 ms of processing) and near-simultaneously (e.g., Pulvermuller et al., 2005; Shtyrov et al., 2005; 2014; Näätanen et al., 2007; MacGregor et al., 2012). Similar rapid brain indices for extracting the meaning and sounds of words have been observed in language production tasks (e.g., Strijkers et al., 2010; 2017; Miozzo et al., 2015; Ries et al., 2017; Janssen et al., 2020; Feng et al., 2021). These results challenge the traditional sequential view on the temporal dynamics of language processing, and instead are better captured by those brain language models which assume words are represented as unified cell assemblies where meaning and sounds ignite rapidly in parallel, both when perceiving (e.g., Pulvermuller, 1999; 2018; Pulvermuller & Fadiga, 2010) and producing words (e.g., Strijkers, 2016; Strijkers & Costa, 2016).

According to the above ‘Integration Models’ (IMs), the neurophysiological data demonstrating sequential dynamics do not concern the first-pass (ignition) activation of language representations. Rather it is assumed to reflect second-pass (reverberatory) activation associated with linguistic operations upon the parallel retrieved words such as verbal working memory, semantic integration, grammatical inflection and articulatory control (e.g. Pulvermuller & Fadiga, 2010; Strijkers & Costa, 2016; Schomers et al., 2017; Strijkers et al., 2017). In other words, IMs predict (at least) two functionally distinct time-courses underlying word processing: Early on the explosion like activation (ignition) of words as a whole indexing target recognition, and afterwards more sequential reverberations linked to task- and context-specific processing. For example, in an influential intracranial study of (written) word production, Sahin and colleagues (2009) observed that depth electrodes in Broca’s region displayed a serial response to the lexical (word frequency) and phonological (phonological inflection of plurals) information during word production planning. The authors interpreted the result as supporting sequential models of language production where lexical information is available well before phonological information. However, within the framework of IMs another interpretation is possible, namely that the later phonological effect did not reflect word form activation per se, but rather reverberatory activation to phonologically inflect the earlier activated target word (i.e., add an ‘s’ to the activated noun form in order to correctly articulate a plural). By incorporating both parallel brain dynamics linked to the neural activation of a word and sequential brain dynamics linked to task- and context-specific linguistic operations upon that word representation, IMs can offer a unified account for the parallel versus sequential time-courses as observed in response to language processing in the literature.

However, proponents of sequential brain language models remain skeptical whether such early parallel activation of lexical and phonological word representations is plausible. One frequently uttered reservation concerns the possibility that these early effects highlight sensorial and/or attentional differences between stimuli-sets rather than linguistic brain activity (e.g., Hagoort, 2008; Mahon & Caramazza, 2008; Indefrey, 2016). Indeed, several of the studies showing that both lexical and phonological brain correlates within the first 200 ms of processing relied on physically different stimuli-sets (e.g., Strijkers et al., 2010; 2017; McGregor et al., 2012; Miozzo et al., 2015). Another criticism is whether ‘fast’ equals ‘parallel’ brain dynamics (e.g., Brehm & Goldrick, 2016; Mahon & Navarrete, 2016). Recent neuromagnetic data taken to support parallel activation of lexical and phonological knowledge in speech production exemplifies this issue: In an MRI-constrained MEG study of overt object naming, Strijkers and colleagues (2017) observed that both word frequency, taken as a metric for lexico-semantic processing, and the place of articulation of the word-initial phoneme (i.e., labial versus coronal), taken as a metric for phonological and phonetic encoding, elicited stimuli-specific fronto-temporal source activity between 160 and 240 ms after stimulus onset. While the authors argued this data pattern best fitted parallel dynamics, the data may still be explained within a sequential framework where functional delays between linguistic components are less than the traditional assumed hundreds of ms (in this example: the lexico-semantic effects occurring in the beginning of the 80ms time-window and the phonological effects emerging near the end of the 80ms time-window). In other words, finding early onsets for distinct word processing components, though requiring the modification of the classical temporal assumptions in sequential models, does not necessarily exclude sequential processing dynamics per se.

To address these criticisms, in the current study we explored the time-course of a lexico-semantic and a phonological variable both in speech production and perception. That is, we directly contrast the same brains, stimuli and, importantly, psycholinguistic manipulations across the two language modalities. Including ‘language modality’ (i.e., production vs. perception) as variable to assess the cortical activation time-course of different linguistic components has a marked advantage compared to prior spatiotemporal research in word production and perception done in isolation: It allows us to assess time-course from a different perspective, which removes both the problem of physical variance and the conceptual problem of whether ‘fast’ equals ‘parallel’. This is because the relevant dimension for testing the hypotheses now becomes the ‘relative’ time-course of the sequence of events *between the language modalities*, rather than the exact, absolute value of when a linguistic component becomes active within each behaviour. That is, regardless of whether a lexico-semantic and phonological manipulation both elicit fast responses in each language modality, what matters is if the time-course of the responses associated with each of the manipulation is different in production and perception. In a sequential model it should since lexico-semantic and phonological activation are functionally separated in the ‘reverse’ order between production and perception. In contrast, in IMs it should not since in both modalities the word, with its lexico-semantic and phonological information, ignites as a whole in parallel. In other words, by exploring whether the time-course of lexico-semantic and phonological word components interacts with production vs. perception, we can assess whether the cortical dynamics of word component activation follows a modality-specific time-course, as predicted by sequential brain language models, or a cross-modal activation time-course, as predicted by parallel models (see Figure 1).

**Figure 1.**
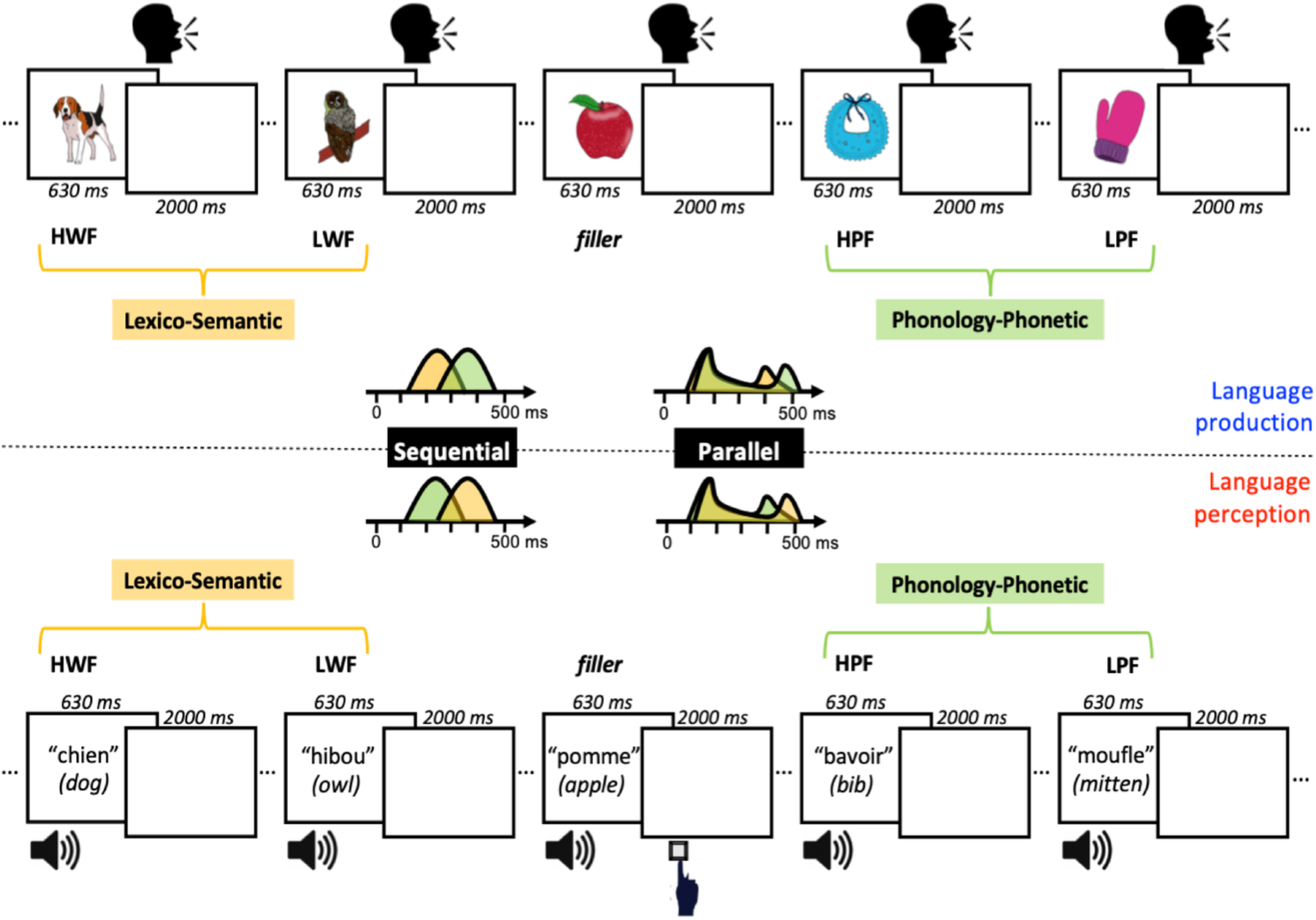
Schematic representation of the experimental design, critical stimuli and hypotheses. In the upper panel an example stream of trial presentations is depicted for language production, where participants have to overtly name the presented objects as quickly as possible. In the lower panel the same example stream of trial presentations is depicted for language perception, where participants listen to spoken words and have to press a response button when a spoken word belongs to the semantic category ‘food’ (go-trials only on filler words; 10% of the trials). For both tasks examples of critical stimuli are depicted, where the contrast between high word frequency (HWF) and low word frequency (LWF) taps into lexico-semantic processing (yellow), and the contrast between high phonotactic frequency (HPF) and low phonotactic frequency (LPF) taps into phonological-phonetic processing (green). Note that in both tasks the exact same stimuli are used. In the middle the temporal predictions for these two variables are schematically depicted where according to sequential models lexico-semantic information (yellow) should be available well before phonological-phonetic information (green) in language production, and the reverse temporal pattern is predicted in language perception. In contrast, according to parallel models lexico-semantic and phonological-phonetic word knowledge should rapidly activate simultaneously both in language production and perception, and only later in the course of processing task-specific differences may emerge between the language modalities.

In order to tap into lexico-semantic and phonological processing, we manipulated two well-known variables which have been successfully applied both in language production and perception. As a metric for lexico-semantic processing, we manipulated word frequency (see Figure 1). The word frequency effect refers to the observation that we are faster in processing words we are well acquainted with (e.g., *dog*) compared to more rare words (e.g., *owl*), and has been taken to reflect the speed of lexical activation in both language production (e.g., Caramazza et al., 2001; Graves et al., 2007; Strijkers et al., 2010; 2011; 2017) and perception (Sereno et al., 1998; Dahan et al., 2001; Dufour et al., 2013; Winsler et al., 2019). As a metric for phonological and phonetic processing, we manipulated word phonotactic probability, which refers to how often phonemes and sequences of phonemes, biphones, of a word co-occur^1^ (see Figure 1). Both in production (e.g., Levelt & Wheeldon, 1994; Vitevitch et al., 2004; Cholin et al., 2006; Laganaro & Alario, 2006) and perception (Vitevitch et al., 1999; Vitevitch & Luce, 1998) it has been found that words with high phonotactic frequency (e.g., horse) are processed more quickly than words with low phonotactic frequency (e.g., glasses); an effect attributed to the speed of word form and/or phonetic processing in both language modalities. This conclusion is further supported by neurophysiological investigations of phonotactic frequency effects displaying differences in a time frame consistent with phonological encoding (e.g., Pyllkanen et al., 2002; McGregor et al., 2012; Dufour et al., 2013; Hunter, 2013; Burki et al., 2015; Den Hollander et al., 2019; Di Liberto et al., 2019).

Relying on these lexico-semantic and phonological indices, respectively, the rationale we followed was to track the ERP responses elicited by these two manipulations, and to compare their time-course within the same participants performing an object naming task, classically used in production, and a go/no-go semantic categorization task which is often used in ERPs studies of perception (Figure 1). With such a design, sequential models predict a specific interaction between ‘language modality’ and the ‘lexico-semantic’ and ‘phonological’ manipulations where the ‘lexico-semantic’ variable should elicit earlier ERP responses than the ‘phonological’ variable for the language production task and the opposite ERP pattern (earlier manifestations of the ‘phonological’ manipulation compared to the ‘lexico-semantic’ manipulation) for the language perception task (Figure 1). In contrast, parallel IMs predict the absence of such interaction since both in the production and perception tasks the ERP responses elicited by the ‘lexico-semantic’ and ‘phonological’ manipulations should emerge simultaneously (Figure 1).

## Methods

In accordance with Open Practices all data (raw and processed), code, analyses pipelines and materials are available in the project OSF repository at https://osf.io/hp2me/.

### Participants

26 participants took part in the experiment (mean age = 21, SD = 3, 20 female). All were recruited through Aix-Marseille University and paid €20 for participation. All participants were right-handed and reported no hearing, speech or neurological disorders, and had normal or corrected-to-normal vision. Two participants were rejected due to equipment failure, and five participants were rejected due to exclusion of a large number of error trials (defined more below), leaving a sample of 19 participants for analysis. The experiment was granted ethical approval by Aix-Marseille Université ‘*Comité Protection des Personnes’ ‘–Committee for the Protection of Individuals’*: CPP 2017-A03614-49.

### Design & Materials

There were two within-participant manipulations of interest, tapping into the lexico-semantic and phonological/phonetic levels of linguistic processing (see Figure 1). The lexical level manipulation was word frequency of the target name (either high frequency or low frequency; mean high log frequency = 1.535, mean low log frequency = 0.5). The phonological/phonetic manipulation was phonotactic frequency of the target name (i.e., summed biphone frequency of words; high frequency = 6836.56, low frequency = 2052.50; and phoneme frequency: high frequency: 254.138, low frequency: 143.122). Word frequency and phonotactic frequency values were retrieved from the Lexique (New et al., 2004) database. The word and phonotactic manipulations were orthogonal, with 220 target items overall. There were thus 110 high word frequency and 110 low word frequency items, and 110 high phonotactic frequency and 110 low phonotactic frequency items (55 items per cell). All stimuli were matched on h-index (number of alternate names given to the image), visual complexity, length of the word in phonemes, bigram frequency, number of phonological neighbors, and duration. Stimuli were significantly different on the crucial word and phonotactic variables (all t-test results show |t| < 2 except the crucial tests; see the stimuli list on the online repository, https://osf.io/hp2me/, for all stimuli, their values on these measures, and the t-test results).

The same target items were used in both the production and perception task, i.e., the stimuli were identical in both tasks. Participants carried out picture naming for the production task and an auditory semantic categorization task for the perception task (a go/no-go task) (see Figure 1). Colored line drawings were retrieved from the MultiPic database (Duñabeitia et al., 2018) and presented in the center of the screen. The auditory stimuli were recorded by a female native French speaker in the soundproof chamber of the *Laboratoire Parole et Langage* of Aix-Marseille University using an RME fireface UC audio interface and a headset Cardioid Condenser Microphone (AKG C520) at a sampling rate of 44 100 Hz. The tokens were segmented and then normalized in intensity to a level of 60 dB using Praat software (Boersma & Weenink, 2020). For the semantic categorization task (perception), participants only responded (gave a ‘go’ response) on filler trials (10% of the trials) which were not analysed (all critical trials had a no-go response). Filler trials consisted of food items, and participants were instructed to press a button if the word they heard was a type of food. There were 25 filler trials leading to 245 trials in each task, with stimuli presented once per task. For the picture naming task (production), participants overtly uttered as quickly and accurately as possible the picture name. Five additional practice trials were present at the beginning of each task, and their data were not recorded.

The stimuli were presented in fully randomised lists per participant and per task. Task order was counterbalanced across participants. The experiment was presented using E-Prime (version 2.0) on a standard computer monitor. Naming responses were collected with a microphone placed in front of the participants for the picture naming task and participants heard the auditory stimuli through non-conductive headphones, and their button press responses on go trials were recorded by a response button box.

### Procedure

Participants first were informed about the study and gave informed consent. Electrodes were fitted to participants (see below) before participants began the experiment proper. Participants were familiarised with the stimuli before the experiment began. For production this entailed seeing all the pictures with their correct name written below, and to be matched in exposure, for perception, this entailed hearing all the stimuli names. Participants began with either the production or the perception task (counterbalanced across participants). During each task participants were able to take a short break (around 10s) every 60 trials, and a longer break (a few minutes) between tasks. After the experiment participants were fully debriefed.

Experiment instructions were written presented in white on a black background and centered in the middle of the screen. Production task trials began with a fixation cross (500ms), followed by a blank screen (500ms), picture presentation (630ms), and ended with an inter-trial interval blank screen (2000ms). Perception task trials followed the same structure and began with a fixation cross (500ms), followed by a blank screen (500ms) before auditory stimulus presentation (approximately 630ms; the average duration of the spoken words), and ended with a blank screen (2000ms) (see Figure 1). The whole experiment lasted 1 hour and 45 minutes with 45 minutes for the experiment proper (including breaks).

### EEG acquisition

EEG was recorded with 64 Biosemi active electrodes with standard 10/20 positioning. Four additional Ag/Ag-Cl electrodes were placed for eye movements (one above and one below the left eye to monitor for vertical eye movements and blinks, two electrodes on the outer canthi of each eye to monitor for horizontal eye movements), and two electrodes were placed on the left and right mastoids for referencing. Electrode impedances were kept below 5 microVolts. The EEG signal was continuously recorded at 512Hz with an online bandpass filter of 0.03Hz – 40Hz using BrainVision software.

### Behavioural data preprocessing

Speech latencies were automatically recorded in E-Prime using a voicekey. Each trial was manually checked (online) to determine if an error was made; errors included production of an incorrect target name, hesitations, and other disfluencies (such as sneezes and coughs). All errors were rejected (14%). In the perception task, correct go-trials (filler items) corresponded to more than 90% performance in all participants; no-go trials (critical items) could of course not be behaviorally analyzed, but in order to keep the symmetry with the production task for the EEG analyses (see below), trials containing errors in the production tasks were also removed from the perception task in each individual (for an average of 14%).

### EEG preprocessing

EEG data, for the production and perception task separately, were preprocessed using FieldTrip (Oostenveld et al., 2011; version 20190716) in Matlab (version R2019a). The raw EEG signal was first filtered with a Hamming windowed sinc filter (one pass, zero phase, cutoff −6dB, maximum passband deviation 0.0022Hz, stopband attenuation −53dB), with a high pass of 0.1Hz (transition width 0.2Hz, stopband 0Hz, passband 0.2-256Hz) and low pass of 40Hz (transition width 10Hz, passband 0-35Hz, stopband 45-256Hz), re-referenced to the average of the mastoids, and epoched from −100ms to 550ms (0ms at target onset). Error data were removed (as explained in *Behavioural data preprocessing*), along with filler trials. Data from each channel was then visually inspected, and channels which were bad were removed and interpolated (using the ‘weighted’ method). Trials with signal above 150mV were also discarded. We then ran an ICA decomposition (with 40 components) to remove components relating to eye movements using the ‘fastica’ method, per participant. Single channels for the horizontal and vertical eye movement channels were created, and each ICA component was correlated with these two channels. All components were visually inspected by plotting their topographies and time courses, and in conjunction with the correlation results artefactual ICA components were rejected before unmixing. Trials were then visually inspected to remove any trials containing excessive noise which had not been captured. If more than 60% trials per condition for each participant were removed during the preprocessing procedure then that participant was rejected from further analysis (five participants were rejected). From the remaining participants, 74% data was retained (i.e., the 14% of trials associated with speech errors as indicated above + an additional 12% of bad EEG signal due to eye movements, etc.).

### Behavioural data analysis

Production latencies corresponding to the cleaned EEG data were analysed. Due to a voice key recording error, latencies for some cleaned trials did not have a latency recorded. After removing these trials, 78% of the production data remained for analysis.

Data were analysed with linear mixed effects models (lme4 package; Bates et al., 2015) in R (R core team 2020; version 4.0.2). Trial was entered as a control variable, and the fixed effects modeled were ‘Word Frequency’, ‘Phonotactic Frequency’, and their interaction. Categorical predictors (the frequency variables) were sum to zero coded, and the continuous predictor (trial) was centred and scaled. Production latency was log-transformed before analysis to reduce skew. The maximal random effects structure that converged contained random intercepts for participant and item (the picture presented), and a random slope of word frequency by participant.

### EEG analysis

In order to determine time windows of analysis in an objective manner (i.e., no ‘visual assessment’ of time-windows), the global field power (GFP) of the baselined EEG signal was calculated (baseline of −100ms to 0ms). GFP is a measurement of the strength of the signal measured across all electrodes, conditions and participants. GFP was calculated per participant per task using FieldTrip in Matlab. GFP was then averaged across participant and task to test for peaks in the GFP signal. GFP peaks were tested for using iPeak (O’Haver, 2020) in Matlab. The reason to explore GFP peaks across tasks (i.e., across production and perception) was to obtain the same time-windows for analyses across the modalities. Using this procedure, three peaks were found at 109ms, 242ms and 340ms (parameters Amplitude threshold = 2.15, Slope threshold = 0.002, Smooth width = 3, Fit width = 6) (see Figure 2).

**Figure 2.**
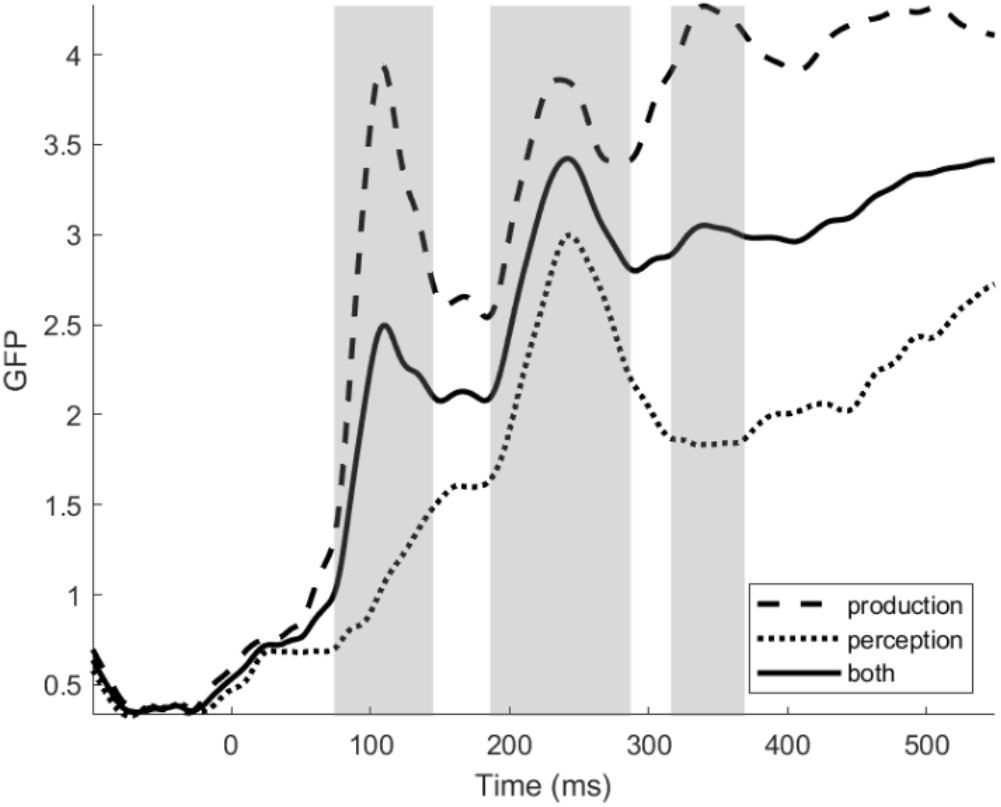
Global Field Power (GFP) to the production and perception tasks in the experiment. GFP was calculated across all electrodes, conditions and participants in production and perception, and the average between the modalities was taken to assess the same time-windows of analyses for production and perception (centered around the maximal peaks). In this manner, three time-windows were objectively determined: 74-145ms, 186-287ms and 316-369ms.

Full-width windows around these peaks were then manually identified. To do this, using the iPeak output per peak, we manually stepped backwards and forwards until reaching the first time point in the backwards or forwards trough. Carrying out the procedure in this way enables asymmetrical widths on either side of the peak, rather than taking a general average of the whole peak window where the peak value may not be centred. This procedure identified the three time windows (TWs) of interest as TW1: 74ms – 145ms, TW2: 186ms – 287ms, and TW3: 316ms – 369ms (see Figure 2). The clean EEG data was then averaged per participant per task per electrode in each of the three time windows, and exported for further analysis.

The data from each time window was tested in R using linear regression and linear mixed effects models (lme4). Channels were recoded by their lateral placement into left (Fp1, AF3, F3, FC3, C3, CP3, P3, PO3, F5, FC5, C5, CP5, P5, AF7, F7, FT7, T7, TP7, P7, PO7, P9, O1), mid (F2, FC2, C2, CP2, P2, F1, FC1, C1, CP1, P1, Iz, Oz, POz, CPz, FPz, AFz, Fz, FCz, Cz), and right (AF4, F4, F6, FC4, FC6, C4, C6, CP4, CP6, P4, P6, PO4, AF8, F8, FT8, T8, TP8, P8, PO8, P10, O2), and recoded by their frontal coronal plane placement into anterior (Fp1, AF7, AF3, F1, F3, F5, F7, Fpz, Fp2, AF8, AF4, AFz, Fz, F2, F4, F6, F8), central (FT7, FC5, FC3, FC1, FT8, FC6, FC4, FC2, FCz, TP7, CP5, CP3, CP1, CPz, TP8, CP6, CP4, CP2, T7, C5, C3, C1, Cz, C2, C4, C6, T8), and posterior (P1, P3, P5, P7, P9, PO7, PO3, POz, Pz, P2, P4, P6, P8, P10, PO8, PO4, O1, O2, Oz, Iz).

For the mixed effects models, all categorical predictors were sum to zero coded, and all continuous predictors were centred and scaled. One model per time window was first run with the main effect predictors Word Frequency (high vs low), Phonotactic Frequency (high vs low), Language Modality (production vs perception), Channel Laterality (left vs mid vs right), and Channel Anteriority (anterior vs central vs posterior)^2^. There were interactions between the channel placement variables, modality, and each frequency variable. Control predictors included trial number, session order (whether participants carried out the production or the perception task first), and average microvoltage in the baseline period (−100ms to 0ms). Due to the size of this model and in order to interpret the interactions (see Results), we then ran models separately for the production and perception data per time window. The control variables were the same as above, and the predictors were Word Frequency, Phonotactic Frequency, Channel Laterality, Channel Anteriority, with interactions between each frequency variable and the two channel variables. The maximum random structure which would converge for all models contained random intercepts for participant and item, and no random slopes. For the linear regression, the same procedure was followed and variables included as with linear mixed models.

For all models t greater than |2| was taken as a marker of significance and confidence intervals were generated using the confint function with the profile method in the lme4 package. P values were generated using the Anova function with type III tests selected in the Car package (Fox & Weisberg, 2019). Post-hoc comparisons were calculated using the emmeans package (Lenth, 2020). To keep the results section focused, only results from predictors of interest and significant effects are reported, but the full results of all models can be found in the supplementary materials and the online repository.

## Results

### Production latencies

Production latencies are listed by frequency in Table 1. High word frequency items were responded to descriptively faster than low word frequency items (Mhigh = 964.81, SD = 368.1; Mlow = 1032.67, SD = 401.2), whereas there was no difference between high and low phonotactic frequency items (Mhigh = 997.67, SD = 391.4; Mlow = 996.89, SD = 379.7).

**Table 1.**
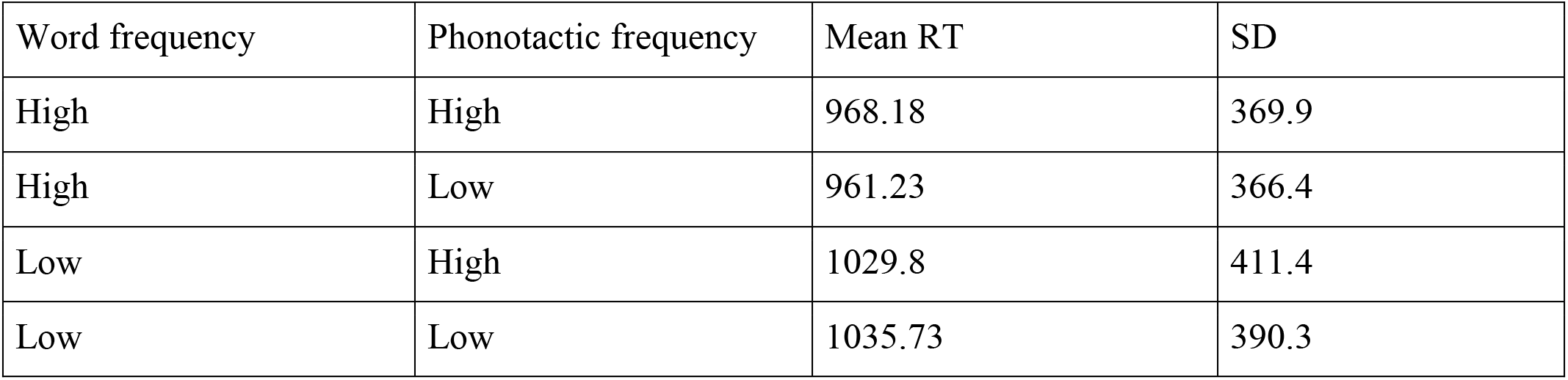
Average production latencies and standard deviation per condition.

The mixed effects models analysis confirmed a significant effect of word frequency (estimate = −0.03, SE = 0.01, t = −2.71, CI = [−0.056, −0.009]), but no significant effect of phonotactic frequency (estimate = −0.0008, SE = 0.01, t = −0.06, CI = [−0.024, 0.022]), nor a significant interaction between the two (estimate = 0.004, SE = 0.01, t = 0.33, CI = [−0.019, 0.027]).

### EEG data: Both modalities

#### Mixed models analysis: TW1 74 to 145ms

In time window 1 there was a main effect of modality (*X^2^*(1,19) > 100, p < 0.001), with lower microvolt recordings in the production task than the perception task. There were no main effects of word frequency nor phonotactic frequency, but phonotactic frequency significantly interacted with modality (*X^2^*(1,19) > 100, p < 0.001). Phonotactic frequency interacted with the channel variable laterality (*X^2^*(2,19) = 12.915, p = 0.002). The three-way interaction between modality, word frequency, and both channel variables was significant (*X^2^*(2,19) = 11.724, p = 0.003; *X^2^*(2,19) = 10.746, p = 0.005), suggesting that the word frequency effect differed in scalp location between modalities. The interaction between modality, phonotactic frequency, and laterality was also significant (*X^2^*(2,19) = 9.812, p = 0.007), suggesting that the phonotactic frequency effect was distributed differently across the scalp in each modality.

In sum, we observed that modality interacted with the word frequency and phonotactic variables; in order to understand the nature of these interactions, separate analyses per modality were performed (see below: *EEG data by modality*).

#### Mixed models analysis: TW2 186 to 287ms

For time window 2, very similar results were found to time window 1. Again, there was a main effect of modality (*X^2^*(1,19) > 100, p < 0.001), and no main effects of word or phonotactic frequency. The word frequency by modality interaction was significant (*X^2^*(1,19) = 17.629, p < 0.001), and the phonotactic frequency by modality interaction was also significant (*X^2^*(1,19) > 100, p < 0.001). The word frequency by anteriority interaction was significant (*X^2^*(2,19) = 7.885, p = 0.020), and the phonotactic frequency by anteriority interaction was significant (*X^2^*(2,19) = 25.072, p < 0.001). The three way interaction between word frequency, modality, and anteriority was significant (*X^2^*(2,19) = 27.915, p < 0.001), and the three way interactions between phonotactic frequency, modality, and both channel variables were also significant (*X^2^*(2,19) = 28.770, p < 0.001; *X^2^*(2,19) = 30.281, p < 0.001). As for TW1, these interactions were further assessed with separate analyses per modality (see below: *EEG data by modality*)

#### Mixed models analysis: TW3 316 to 369ms

In TW3, the same main effects patterns were found: main effect of modality (*X^2^*(1,19) > 100, p < 0.001) and no main effects of word and phonotactic frequency. There was a word frequency by modality interaction (*X^2^*(1,19) = 13.654, p < 0.001), and word frequency by anteriority interaction (*X^2^*(2,19) = 16.552, p < 0.001). The three way interaction between word frequency, modality, and each channel variable were significant (*X^2^*(2,19) = 35.459, p < 0.001; *X^2^*(2,19) = 6.346, p = 0.042). Phonotactic frequency was only involved in a three way interaction, with modality and anteriority (*X^2^*(2,19) = 10.663, p = 0.005). Separate analyses per modality were performed in order to assess the nature of these interactions (see below: *EEG data by modality*).

### EEG data: by modality

#### Mixed models analysis: TW1 74 to 145ms

##### Production

Main effects of word frequency and phonotactic frequency were not significant. Word frequency however interacted with laterality (*X^2^*(2,19) = 11.90, p = 0.003) (see Figures 3 and 4). Post-hoc t-tests revealed that this interaction was driven by a significant increase from right to left electrodes for low frequency items (p < 0.001), which was absent for high frequency items (p = 0.789). There was also a significant interaction between phonotactic frequency and laterality (*X^2^*(2,19) = 6.974, p = .031) (see Figures 3 and 4). Post-hoc tests showed this interaction was driven by a marginally significant difference between high and low frequency items at left electrodes (p = 0.056), and a significant increase from right to left electrodes for high frequency items (p < 0.001), which was absent for low frequency items (p = 0.998).

**Figure 3.**
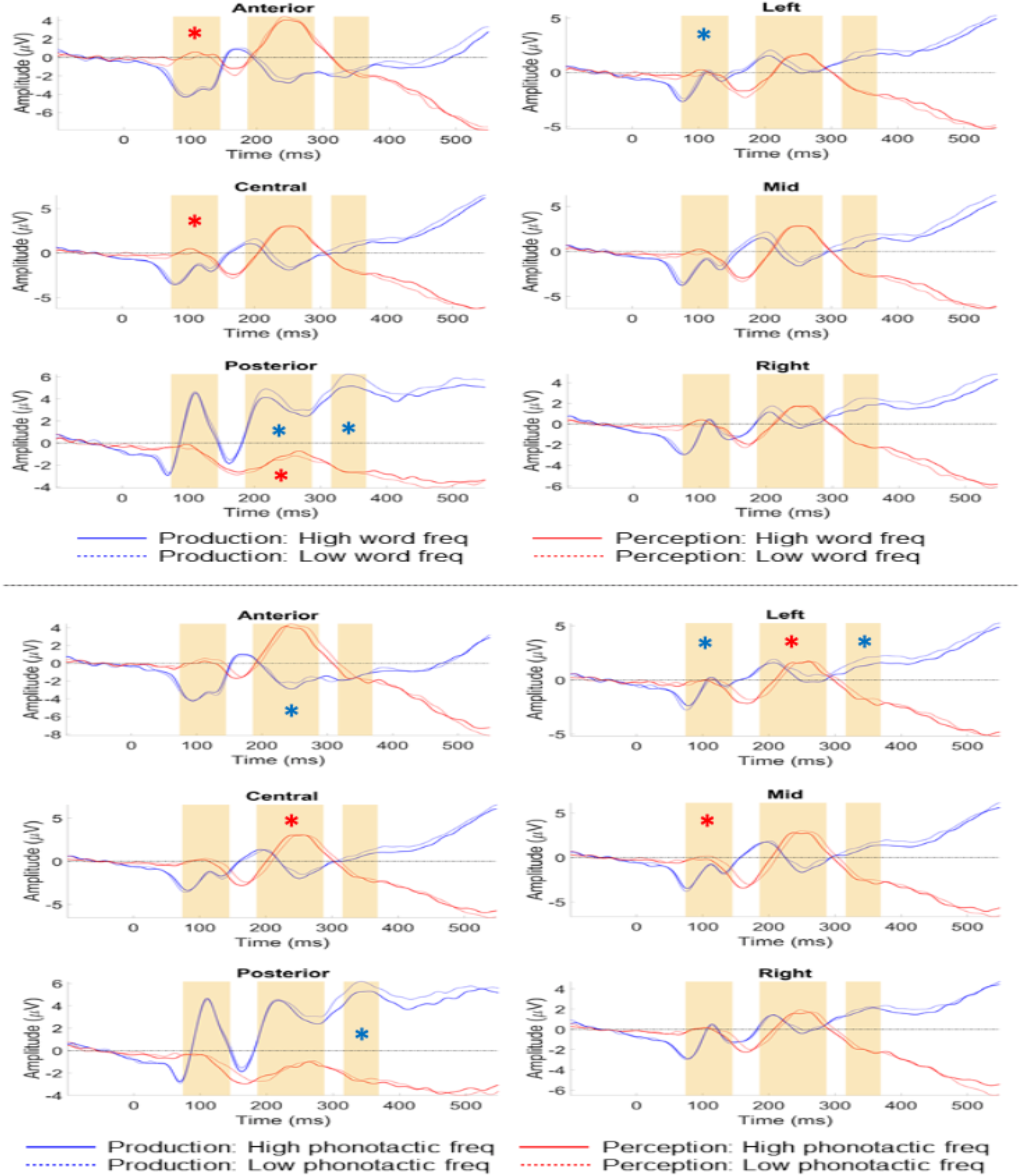
ERP results for the production and perception tasks at each electrode location. The upper panel displays the ERPs for high (full line) versus low (dotted line) word frequency in production (blue) and perception (red). The lower panel displays the ERPs for high (full line) versus low (dotted line) phonotactic frequency in production (blue) and perception (red). Time-windows for analyses are highlighted with yellow rectangles, and the significant interactions of scalp site with word or phonotactic frequency, respectively, are depicted with a blue asterisk for production and a red one for perception. Note that in production effects for both variables were significant in all three time windows, while in perception effects for both variables were significant for the first two time windows, but not for the last time window. Note that for visualisation purposes ERPs are presented here as the averaged means, but the actual data-analyses were performed for single-trial ERP data with mixed linear effect models.

**Figure 4.**
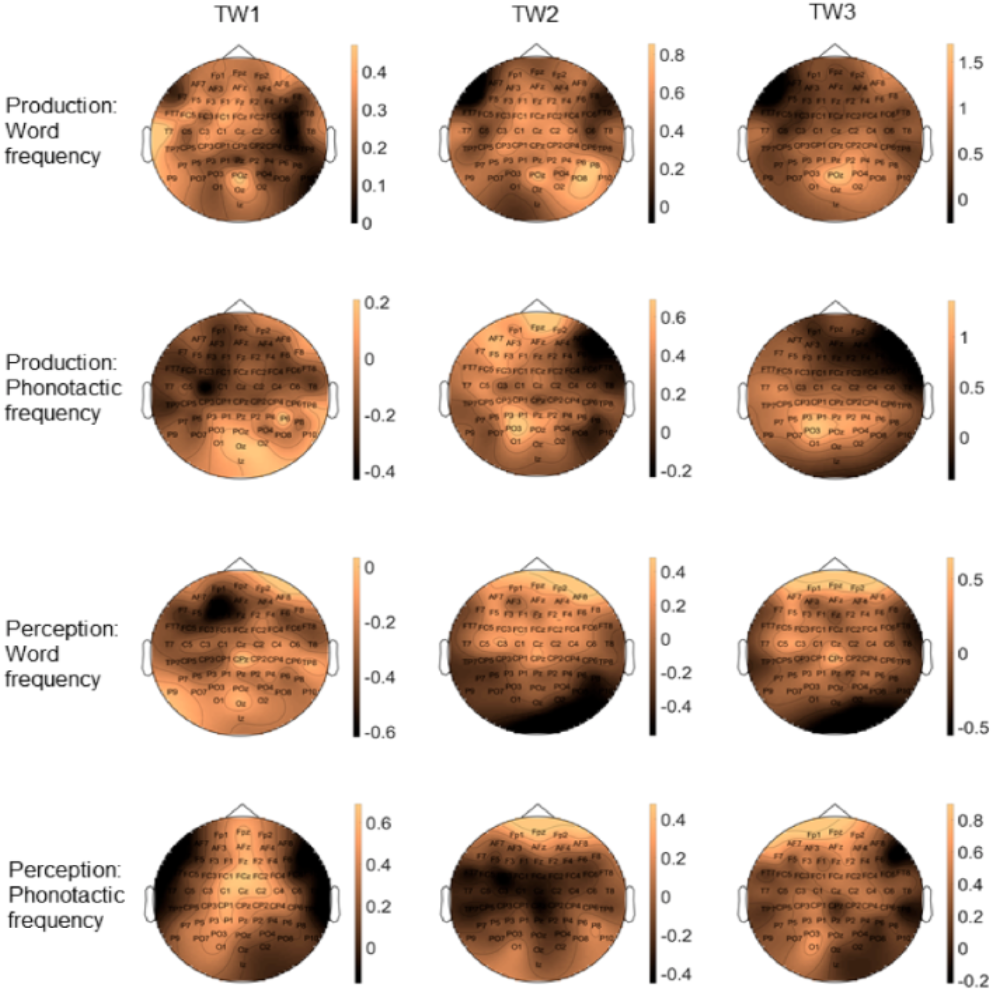
Topographic scalp distributions of the word frequency and phonotactic frequency effects in production and perception at each time window of analyses.

##### Perception

Similarly to production, there were no main effects of word or phonotactic frequency. There was however a significant interaction between word frequency and anteriority (*X^2^*(2,19) = 11.419, p = 0.003) (see Figures 3 and 4). Post-hoc tests suggest this interaction is due to high word frequency items being more positive than low frequency items across the scalp, with a greater voltage increase at anterior sites (high frequency, p < 0.001; low frequency, p = 0.045) and central sites (high frequency, p = 0.086; low frequency, p = 0.700) compared to posterior sites. For phonotactic frequency we found a significant interaction with laterality (*X^2^*(2,19) = 16.487, p < 0.001), with a trend towards significance between low and high frequency items at mid electrodes (p = 0.064), a significant voltage increase from left towards mid sites for low frequency items (p < 0.001), which was absent for high frequency items (p = 0.988), and a significant increase from mid to right sites for high frequency items (p < 0.001), which was absent for low frequency items (p = 0.610) (see Figures 3 and 4).

#### Mixed models analysis: TW2 186 to 287ms

##### Production

There were no main effects of word nor phonotactic frequency. Word frequency did interact with anteriority (*X^2^*(2,19) = 7.477, p = 0.024) (see Figures 3 and 4). Post-hoc tests showed a stronger increase in positivity from anterior to posterior electrodes for low frequency items (p < 0.001; diff. = 1.0mV) compared to high frequency items (p < 0.001; diff. = 0.6mV). Phonotactic frequency also interacted with anteriority (*X^2^*(2,19) = 41.151, p < 0.001), with a marginally significant difference between high and low frequency items at anterior sites (p = 0.059), and a stronger increase in positivity from central to anterior sites for low frequency items (p < 0.001; diff. = 1.0mV) compared to high frequency items (p < 0.001; diff. = 0.5mV) (see Figures 3 and 4).

##### Perception

No main effects of word or phonotactic frequency were found. The interaction between lexical frequency and anteriority was significant (*X^2^*(2,19) = 30.431, p < 0.001), where post-hoc inspection suggests a stronger increase in negativity from anterior to posterior electrodes for low frequency items (p < 0.001; diff. = 3.7mV) compared to high frequency items (p < 0.001; diff. = 3.2 mV) (see Figures 3 and 4). Also the phonotactic frequency by anteriority interaction was significant (*X^2^*(2,19) = 14.842, p < 0.001), with a trend for low frequency items being more negative than high frequency items at central electrodes (p = 0.069), and a stronger increase in positivity from posterior to central sites for high frequency (p < 0.001; diff. = 2.8mV) then low frequency items (p < 0.001; diff. = 2.4mV). The phonotactic frequency by laterality interaction was also significant (*X^2^*(2,19) = 32.178, p < 0.001), with trends for low frequency items being more negative than high frequency items at left (p = 0.08) and mid sites (p = 0.100), and a stronger increase in negativity from right to left electrodes for low frequency items (p < 0.001; diff. = 0.5mV) compared to high frequency items (p = 0.030; diff. = 0.1mV) (see Figures 3 and 4).

#### Mixed models analysis: TW3 316 to 369ms

##### Production

No main effects of either frequency variable were found. Word frequency interacted with anteriority (*X^2^*(2,19) = 50.954, p < 0.001) (see Figures 3 and 4). Post-hoc tests demonstrated a trend towards more positivity for low compared to high frequency items at posterior sites (p = 0.068), and a stronger increase from frontal towards posterior electrodes for low frequency (p < 0.001; diff. = 2.4mV) then high frequency items (p < 0.001; diff. = 1.8mV). Phonotactic frequency interacted with both anteriority (*X^2^*(2,19) = 11.452, p = 0.003) and laterality (*X^2^*(2,19) = 6.458, p = 0.040) (see Figures 3 and 4). Post-hoc tests suggest that the phonotactic frequency by anteriority interaction was mainly driven by a stronger increase in positivity from anterior to posterior sites for low (p < 0.001; diff. = 2.2mV) compared to high frequency items (p < 0.001; diff. = 1.8mV), and the phonotactic frequency by laterality interaction was driven by the presence of a significant decrease in positivity from mid to left sites for high frequency items (p = 0.03), which was absent for low frequency items (p = 0.967).

##### Perception

No main effects of word frequency or phonotactic frequency were found, nor any interactions with the channel variables (see Figures 3 and 4).

#### Additional analyses: Linear regression

We also carried out a linear regression analysis on the three time windows, akin to a more typical ANOVA style analysis of the data, given that this still concerns the most performed analyses on neurophysiological data and is therefore interesting as a comparison. For brevity and clarity, we will not discuss the statistical details of these analyses - they can be consulted in full in the online repository at https://osf.io/hp2me/ - but only descriptively report the main findings. In general, the results were quite similar to the mixed models analysis in that they confirmed the presence of word and phonotactic frequency effects for both modalities in all time windows (with the exception of TW3 in perception; just as in the mixed models analyses). The biggest, and quite noteworthy, difference with the mixed models analyses is that the linear regression analyses displayed main effects of both frequency variables in almost all analyses. In other words, the word and phonotactic frequency effects were much more marked and broader in the traditional regression analyses compared to the mixed effect models analyses.

A deeper investigation into these discrepancies suggested that there is a substantial amount of variance attributable to the items used. This variance is accounted for as random variation in the mixed effects models, i.e., the variance is treated as ‘noise’ and not as part of the variables of interest. In linear regression, this variance is instead (at least partially) grouped with variation in the variables of interest, leading to a higher chance of a significant effect. In this manner, we believe that the mixed effect models analyses is a better approach to single out the “pure” frequency effects, since item variance between conditions is better controlled for compared to traditional ANOVAs and linear regression analyses. This difference between the types of analyses might have important consequences for all neurophysiological research which relies on conditions containing physically different stimuli. For present purposes, we will not go any further into this interesting methodological issue (but see Fairs et al., 2020), and instead highlight that both analyses point to the same theoretical insight, namely early parallel access of word and phonotactic frequency in both production and perception. Below we will discuss what this finding signifies for the temporal dynamics of brain language models.

## General Discussion

We investigated the temporal dynamics of word activation by comparing the event-related brain potential elicited by lexico-semantic and phonological knowledge in production versus perception, and analysing the data at the single-trial level with mixed linear effect models. The results showed significant word frequency and phonotactic frequency effects (in interaction with scalp location, respectively) in early time windows (74-145ms and 186-287ms) for both word production and perception (see Figure 3). At a later time window (316-369ms) word and phonotactic frequency effects only remained significant for word production, but not for word perception (see Figure 3). This data pattern provides evidence for ultra-rapid parallel activation dynamics underpinning word retrieval in both production and perception. The speed of linguistic activation, it’s parallel nature for different word knowledge and the identical temporal profile (early on) when speaking and understanding, places important novel constraints on brain language models: (1) The data provide strong evidence for parallel IMs where words are represented in the brain as unified cell assemblies across the language modalities (e.g., Pulvermuller, 1999; 2018; Strijkers, 2016; Strijkers & Costa, 2016); (2) The early parallel activation which is followed at a later stage of processing by modality-specific activation fits the predictions of Hebbian-like neural assembly models where early word ignition is identical between production and perception and later reverberation is task- and context-specific; (3) The similarities in cortical dynamics between word production and word perception suggests the same mental representations underpin both modalities and therefore production-perception interactions in conversation likely tap into the same neural resources for lexical representations between speakers and listeners.

### (1) Parallel cortical dynamics of word processing

The main objective of this study was to shed novel light on a central and longstanding question in the psycho- and neurolinguistic literature: Are words processed in a hierarchical sequential manner or in a parallel integrated manner? The data were clear-cut and demonstrated that lexico-semantic and phonological-phonetic word knowledge are rapidly activated in parallel, both when uttering and comprehending words. The observation of an identical activation time-course across the language modalities is crucial, since the parallel effects cannot be attributed to physical variance between stimuli nor to a cascade of activation over sequentially organised levels of word processing.

With regard to physical variance between stimuli (e.g., sensorial differences between low and high frequency stimuli), in the present study the input is different between production (visually presented objects) and perception (auditory presented words), but nonetheless the same temporal effects between the modalities were present for the word frequency and phonotactic frequency effects. Therefore, explaining the data as due to sensorial differences between the low and high frequency conditions is highly unlikely (if not impossible) since it would mean that by chance the same sensorial differences are present for both the visual and auditory input, for both manipulations and in both early time-windows. Instead, parsimony clearly favors a linguistic explanation since across the two modalities identical word processing components are targeted. Adding further support to this conclusion is the observation that we replicated the typical ERP morphology found for word frequency and phonotactic frequency in prior production and perception studies in isolation. In production ERPs elicited by low frequency items (both word and phonotactic) display more positive going waveforms than high frequency items (e.g., Strijkers et al., 2010; 2011; Baus et al., 2014; Burki et al., 2015), while in perception ERPs elicited by low frequency items typically display more negative going waveforms than high frequency items (e.g., Hauk et al., 2006; McGregor et al., 2012; Hunter, 2013; Winsler et al., 2019). Here both these ERP patterns were replicated and integrated within the same participants and for the same stimuli, lending strong reliability to the observed results.

The current cross-modal timing cannot be reconciled with those brain language models assuming functional delays between word components in the range of 100s ms (e.g., Friederici, 2002; 2011; Indefrey & Levelt, 2004; Indefrey, 2011). Moreover, the argument that the overlapping time-course of lexico-semantic and phonological word knowledge could reflect fast sequentially and interactivity rather than parallel activation finds little support in the current results. If lexico-semantic and phonological-phonetic word processing are functionally (and thus temporally) distinct, we should have seen an interaction between the language modalities with phonotactic frequency manifesting *prior* to word frequency in perception *and* that same phonotactic frequency manifesting *after* word frequency in production. No such interaction was present, and instead the ERP data revealed effects of lexico-semantic and phonological word processing which were indistinguishable between language production and perception in our early time-windows (74-145ms; 186-287ms).

The fact that we can contrast the activation time course of a lexico-semantic and phonologic variable across two modalities, offers an important advantage over the traditional within-modality studies, namely that the points of comparison are doubled. For example, taking the first time window where we observed significant effects for both variables (74-145 ms), this means that in order to claim fast sequentiality the maximal possible onset delay that remains between lexico-semantic and phonological access in production and (vice versa respectively) in perception has to be less than 35 ms. At this point one may question how such small (potential) delays would still support the notion of sequentiality. For one, these short delays perfectly fit with axonal conduction velocities between different brain regions (e.g., Miller, 1996; Matsumoto et al., 2007), hereby denoting physical distance delays in the brain, not functional ones. In a similar vein, it has been recently shown from recordings with micro-wire electrodes that small spatiotemporal delays around 30ms offer an ideal temporal window in the human brain to form Hebbian cell assemblies (e.g., Roux et al., 2021). Furthermore, if sequential activation would be so fast that it becomes indistinguishable to asses between such early time-windows (as observed here), what happens for the remaining 400-500 ms in those models? The elegance of sequential models is that they link different linguistic operations to different points in time that gradually enroll over the entire period of speech planning. Unlike for parallel IMs (see also next section), no theory nor predictions are available for sequential models which would explain what happens during the bulk of speech planning if the serial activation onsets are as fast as observed here. Also, assuming rapid interactivity does not explain the data pattern we observed in the present study: In interactive models (e.g., Dell, 1986; Friederici, 2011) lexico-semantic and phonological processing can overlap due to the feedback activity from a layer higher in the hierarchy (e.g., phonology) onto a layer lower in the hierarchy (e.g., lexical). However, in this case the overlap produces itself after initial sequential feedforward activation. In other words, an interactive brain language model predicts first sequential activation dynamics followed by parallel activation dynamics, while in the current study we observed the reverse pattern with parallel effects very early on and modality-specific effects later on. In sum, for sequential and interactive models to account for the current data pattern, critical properties why hierarchical sequentiality or interactivity was proposed as temporal processing dynamic to begin with would need to be entirely dropped or changed. In contrast, parallel IMs exactly predicted the temporal pattern we observed in this study, even the ultra-rapid parallel effects in the earliest time-window (74-145ms) (e.g., Pulvermuller, 2018).

While in language perception a few of such very early effects linked to word processing have been reported before (e.g., Pulvermuller et al., 2005; McGregor et al., 2012; Grisoni et al., 2016; Strijkers et al., 2019), to our best knowledge this is the first demonstration of ultra-rapid word access in language production. Although we do not believe the early effect indexes item-specific retrieval of the target word, for which the time-window between 186ms and 287ms seems a better candidate based on the previous literature (e.g., Vihla et al., 2006; Costa et al., 2009; Strijkers et al., 2010; 2017; Aristei et al., 2011; Miozzo et al., 2015; Python et al., 2018; Feng et al., 2021), its presence both in the production and perception tasks does suggest a linguistic effect. We argue this early effect reflects a difference of “global” word activation between the conditions, with high word and phonotactic frequency items causing enhanced activation of a set of potential words linked to the input compared to the low word and phonotactic frequency items. In other words, the differences between the high and low frequency conditions trigger almost immediately upon perceiving the input activation within the linguistic system which is different in its magnitude between frequent and more rare words (this difference could be due to higher resting level of activation and/or stronger connectivity and myelinization between input features and lexical features for the more frequent words). Note also that such ultra-rapid linguistic activations fit well with recent spatiotemporal evidence suggesting that combinatorial processes emerge already within 200 ms of processing (e.g., Pylkannen, 2019). Clearly, if we can start integrating two words at such speeds, it logically follows that some information of each single word has to be retrieved even faster.

In sum, the uncovered parallel cortical dynamics of distinct word components in both production and perception cannot be accounted for by the dominant sequential hierarchical brain language models in the literature (e.g., Friederici, 2002; 2011; Indefrey & Levelt, 2004; Hickok & Poeppel, 2007; Indefrey, 2011; Hickok, 2012). In contrast, the data nicely fit parallel IMs which state that words rapidly ignite as whole in parallel, and do so identically in language production and perception (e.g., Pulvermuller, 1999; 2018; Strijkers, 2016; Strijkers & Costa, 2016). To our best knowledge, this is the strongest demonstration yet for parallel cortical dynamics underpinning word processing and the main contribution of our study.

### (2) Ignition and reverberation as neurophysiological principles of word processing?

Another intriguing observation in our data was that for the late time-window of analysis (316ms - 369ms) language production and perception displayed different patterns. Both the word and phonotactic frequency effects remained significant for the language production task, but were no longer significant for the language perception task^3^. This pattern, where early on in the course of processing the cortical dynamics are the same between production and perception, but later on they become different, fits with the theoretical assumptions put forward in Hebbian-based neural assembly models of word processing (e.g., Pulvermuller, 1999; 2018; Strijkers, 2016; Strijkers & Costa, 2016). Within such models, the idea is that early ignition of a cell assembly and later reverberation within that cell assembly can fulfil distinct functional roles. The notion stems from the neurophysiological properties of cell firing and theoretical systems neuroscience (e.g., Hebb, 1949; Braitenberg, 1978; Fuster, 2003; Buszaki, 2010; Singer, 2013). The firing of neuronal cells is marked by a rapid, explosion-like onset (ignition), followed by slower, sustained activity (reverberation). Importantly, while originally it was assumed this slow reverberatory decay of neural activity after cell ignition was mainly noise in the system, by now it is clear that reverberation is sensitive to stimuli- and task-relevant properties (e.g., Fuster, 1995; Singer & Gray, 1995; Tsodyks & Markram, 1997; Azouz & Gray, 2003). In this manner, cell ignition has been linked to ‘target identification’, and reverberations upon the ignited cells to second-order processes such as memory consolidation, reprocessing, decision-making and meta-cognition (e.g., Mesulam, 1990; Fuster, 1995; 2003; Engel et al., 2001; Dehaene et al., 2006).

A similar idea of distinct functional states linked to the different neuronal firing properties of ignition and reverberation can be adopted to language processing (e.g., Pulvermuller, 1999; 2002; 2018; Strijkers, 2016; Strijkers & Costa, 2016; Strijkers et al., 2017), and its plausibility has been computationally demonstrated (e.g., Garagnani et al., 2008; Schomers et al., 2017). Specifically, for word processing, the hypothesis is that rapid ignition would reflect the activation of the memory representation of a word associated with the input, and reverberation with reactivation of the ignited word assembly linked to task- and behaviour-specific processing such as verbal working memory, semantic integration, grammatical inflections and articulatory control (e.g., Pulvermuller, 2002; Strijkers, 2016; Strijkers et al., 2017). For example, if one intends to utter the phrase ‘(s/*he) runs*’, during ignition the “base” representation ‘*(to) run*’ would quickly become activated, and subsequently during reverberation this “base” representation is morphologically inflected to ‘*runs*’ in function of the syntactic context and constraints (*s/he*). So how could such neurophysiological dynamics of ignition and reverberation account for the fact that in our data word and phonotactic frequency effects remained significant for the language production task, but not for the perception task? A possible answer lies in the nature of the task and behaviour that participants had to produce: In production, the goal-directed behaviour of the participants is to utter the specific word associated with the input. Hence, the input-specific word representation, with its associated lexico-semantic and phonetic knowledge, has to be kept online in order to successfully execute the task and articulate the unique word response. In contrast, in the perception task, the goal-directed behaviour of the participants is geared towards the assessment whether the word associated with the input fits in the general category knowledge of ‘food-items’. In other words, in the production task the focus is on the single word representation that needs to be uttered without activating other potential word candidates that could interfere with the goal-directed behaviour. However, in the perception task, the activated word representation needs to be integrated with other word and category knowledge. This concurrent activation unrelated to the unique item conveyed by the input may therefore cause less pronounced word-specific effects in the perception task compared the production task (see also footnote 3).

From this point of view, the full electrophysiological data pattern we observed in our study can be accounted by the neural assembly model of words in the brain in the following manner (see Figure 5): After early initial sensorial activation in response to the input, (1) the first linguistic activation emerges roughly between 75-150 ms after onset denoting the start of “globally” activating potential words fitting the initial sensory analyses, with frequent words triggering a larger set of potential candidates than less frequent words; (2) next, this initial “global word space” activation is further refined and delineated within roughly 150-250 ms leading to the ignition and recognition of the specific lexical item associated with the input; (3) finally, roughly after 300 ms, slower, sequential-like reverberation upon the activated word assembly takes effect in order to embed the recognised target word into the proper linguistic and task context to be able to perform the intended behaviour, and modality-specific processing effects become visible. Note that within this framework the sequential effects of word processing reported in the literature (and discussed in the Introduction; e.g., Indefrey & Levelt, 2004; Friederici, 2011; Indefrey, 2011) would thus stem from reverbatory processing, but not ignition. Put differently, parallel neural assembly models do not negate the presence and even necessity of serial-like operations in word production and perception, but rather link those observations to a different functional interpretation. That said, the current study does not prove this reverbatory link per se and the proposed framework is tentative. Nevertheless, given the same early parallel effects between production and perception as proposed by parallel IMs, and the prediction of these models that later on in the processing sequence task-specific differences will emerge between production and perception, makes this interpretation a relevant one. Future investigations into this framework of functionally distinct linguistic operations associated with cell ignition and reverberation are important, since it may provide us with linking hypotheses between the algorithms of language processing and their neural implementation.

**Figure 5.**
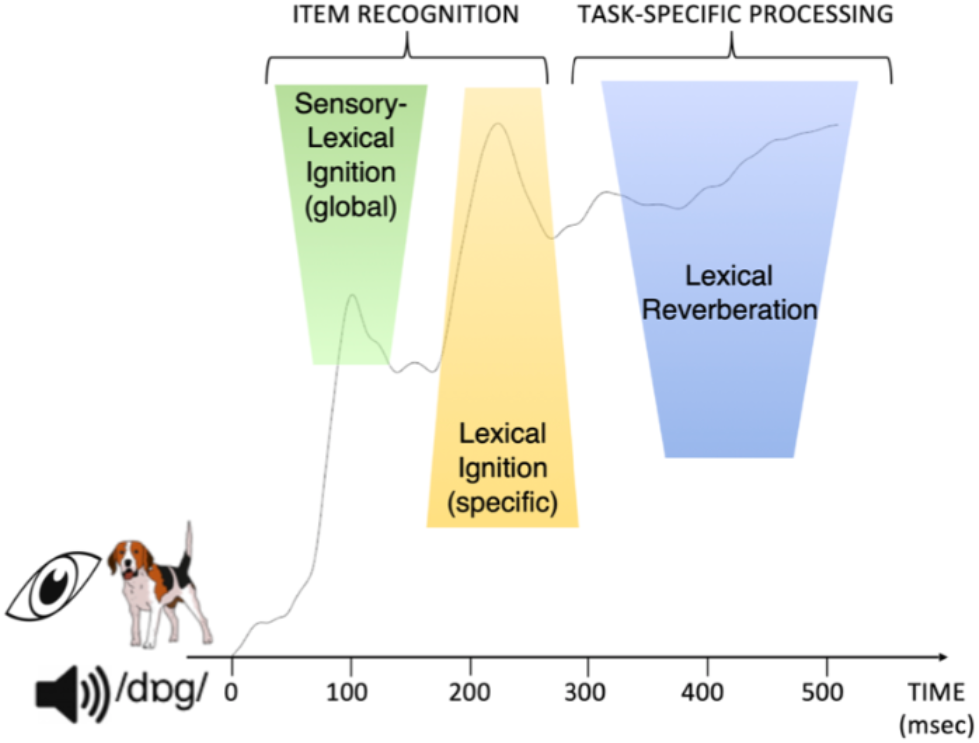
Schematic overview of word processing in a neural assembly framework of ignition and reverberation. Depicted is the global field potential elicited by speech production and speech perception. (1) Upon sensory analyses of a visual stimulus (e.g., picture or spoken word ‘dog’), the first signs of lexical activation occur very early on (from 75ms onwards) during ‘global’ sensory-lexical ignition where potential lexical candidates associated with the input become partially activated; (2) From 175ms onwards lexical ignition takes effect where the word assembly corresponding to the input becomes active and selected for further processing; (3) From 300ms onwards lexical reverberation manifests where the word assembly is re-activated (fully or partially) in function of the specific task and behavior.

### (3) A blueprint of perception-production interactions for conversational dynamics

To conclude, our data are also relevant to constrain integrated models of production-perception interactions and conversation. While traditionally both language modalities were investigated separately, more recently, there is increased interest about the nature of language processing under more ecologically valid conditions such as the dyad (e.g., Hasson et al., 2012; Pickering & Garrod, 2013; Schoot et al., 2016). Importantly, investigations contrasting speaker-listener dynamics, have demonstrated neural overlap between the modalities for high-level linguistic operations such as syntactic unification of message-level information (e.g., Menenti et al., 2011; Segaert et al., 2012; but see Matchin & Wood, 2020), or communicative alignment in conversation (e.g., Stephens et al., 2010; Silbert et al., 2014; Dikker et al., 2017). These data highlight that the language production and perception systems are more shared than originally thought. Less evident to distil from these data is whether the encoding and decoding of language recruit identical neural populations or rather two interactive, but segregated neural populations which are integrated at the highest levels of linguistic processing (e.g., communicative common ground). Indeed, that processes such as communicative alignment or syntactic unification tap into the same neurobiological substrates for production and perception does not necessitate the assumption that lower-level building blocks equally rely on shared representations across modalities (e.g., Hagoort & Indefrey, 2014; Pickering & Garrod, 2014; Silbert et al., 2014), and most brain language models still assume at least partial separation between production and perception at the level of words (e.g., Scott & Johnsrude, 2003; Indefrey & Levelt, 2004; Hickok & Poeppel, 2007; Friederici, 2011; Hickok, 2012; Hagoort & Indefrey, 2014). Addressing whether this is indeed the case is an important endeavor for future research in order to mechanistically link high-level linguistic processing between interlocutors with the basic linguistic elements on which they rely.

The current data may provide some hints towards this objective, given the overlapping activation time-course between production and perception at the level of words. This temporal dynamic fits the notion that words, and their respective linguistic components, share the same cortical representations in production and perception. And though it remains plausible that the time-course of processing may be similar between perception and production, but their spatial and functional arrangement is not - an important question to be explored in future research - the similar ERP pattern across the language modalities as obsered here neatly follows the predictions of IMs (e.g., Pulvermuller, 1999; 2018; Strijkers, 2016; Strijkers & Costa, 2016). Key to neural assembly IMs is that representations are formed through Hebbian-like learning binding together correlated input (perception) - output (production) associations (e.g., Pulvermuller & Fadiga, 2010; see also Skipper et al., 2017; Shamma et al., 2021); this way, the mental representations of words rely on distributed action-perception circuits which integrate traditional “production properties” (e.g., motor knowledge) and “perception properties” (e.g., acoustics) into a single coherent unit. From this perspective what differentiates language production and perception are not the linguistic representations, but the dynamics upon those identical representations. If so, this will have important consequences for interweaving production and perception.

For example, a key element in current conversational models is the assumption that successful alignment (i.e., the coordination and copying of utterances between interlocutors) relies on predictive processing (e.g., Federmeier, 2007; Pickering & Garrod, 2013; 2014; Dell & Chang, 2014; Silbert et al., 2014). In short, a prominent idea is that during perception we rely on the production system to make predictions about the upcoming sensorial information and vice versa during production we rely on ‘efference copies’ in the perception system (e.g., Pickering & Garrod, 2007; 2013; Scott et al., 2009; Tourville & Guenther, 2011; Martin et al., 2018). For this mechanism to work we thus need two separate (though closely interacting) representations, one in production and one in perception. However, if production and perception rely on the same neural representations for words, then this matching mechanism with predictions in one modality and implementations in the other modality wouldn’t work. The latter doesn’t mean that prediction wouldn’t play a role in conversation; it likely does, and in fact, the very early ERP responses to word knowledge we observed here (75-150 ms) seem a prime candidate to reflect such “partial” predictions of possible words (e.g., Grisoni et al., 2016; Strijkers et al., 2019). But the mechanism to implement that prediction would need to be different, where at least the top-down prediction and bottom-up driven analyses of the input converge on the ‘same’ neural representation. Similarly, for speech error monitoring, which necessitates a close interplay between production and perception (e.g., Hickok, 2012; Pickering & Garrod, 2013), the current data fit well with recent findings demonstrating that forward predictions stemming from the cerebellum go beyond mere articulatory control, but include lexical and phonological levels of processing (e.g., Runnqvist et al., 2016; 2020). That is, if words are reflected in the brain as holistic action-perception cricuits, it follows that sensory-motor to cerebellar connectivity will also affect more abstract levels of linguistic processing. Finally, and still largely uncharted territory, the manner to link up low-level language processes with higher-level ones in conversational contexts will be markedly different whether the same or different word representations need to be unified onto a common ground between speakers and listeners. Continued research exploring high-level production-perception interactions in ecologically valid conversational contexts, together with more basic, controlled investigations of low-level spatiotemporal similarities and differences between production and perception will be valuable endeavors to address these questions.

## Conclusion

Our ERP data revealed surprisingly rapid and parallel activation of lexico-semantic and phonology-phonetic word knowledge, which was indistinguishable between the production and perception of language. These data contrast with the traditional sequential hierarchical models of word production and perception, and instead offer strong support for parallel integration models based on Hebbian-like cell assembly theory where words reflect the Gestalts of language in the brain (e.g., Pulvermuller, 1999; 2018; Strijkers, 2016; Strijkers & Costa, 2016). Additional support for this conclusion stems from the observation that the early cross-modal parallel activation of word components was followed later on by distinct ERP effects between production and perception. We interpreted these findings within a neural framework of cell ignition and reverberation being associated with different cognitive states of word processing: rapid, parallel ignition with word activation and retrieval, and later reverberation with behaviour-specific processing upon the ignited word assembly in function of language modality. In this manner, this study may offer a novel and exciting way to map basic neurophysiological brain dynamics onto cognitive operations necessary to express linguistic behaviour.

## Author contributions

K.S. designed to study; K.S., A.M. and S.D. implemented the experiment and collected the data; A.F. analysed the data; A.F. and K.S. interpreted the data and wrote the manuscript; all authors reviewed the manuscript.

## Acknowledgements

This work is financed and supported by a research grant of the ‘Agence National de la Recherche’ (ANR) awarded to the last and corresponding author in this study (Kristof Strijkers): ANR-16-CE28-0007-01. Furthermore, the study received additional support by two ANR grants awarded to the Institute of Language, Communication and Brain (ILCB; ANR-16-CONV-0002) and the Brain Language Research Institute (BLRI; ANR-11-LABX-0036).

## Footnotes

1 Note that phonotactic frequency and phonotactic probability are often used interchangeably in the literature (e.g., Vitevitch et al., 1999); here we prefer the term phonotactic frequency because our measure focusses on the summed biphone frequency, and not segment probability (a measure sometimes used in combination with biphone frequency for phonotactic probability).

2 Model in R syntax for ERP analyses (example here for TW1): 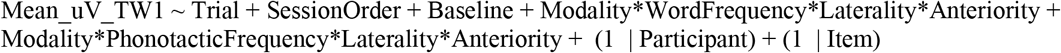

3 Note, when inspecting visually the figures (see Figure 3), it seems that after TW3 word and phonotactic frequency effects become significant again in perception as well. To assess this, post-hoc we performed an additional mixed linear effect analyses between 400 and 500 ms after stimulus onset. The results of these post-hoc analyses (which can be consulted in detail in the online repository: https://osf.io/hp2me/) indeed confirmed significant interactions of both word and phonotactic frequency with electrode location in perception (and also in production). Nevertheless, this post-hoc result does not change the fact that, in contrast to the early time-windows, from around 300 ms after stimulus onset the ERP effects are different between production and perception, with less robust and extended word and phonotactic frequency effects in perception compared to production.

